# Obesity-induced gut microbiota dysbiosis can be ameliorated by fecal microbiota transplantation: a multiomics approach

**DOI:** 10.1101/654228

**Authors:** Maria Guirro, Andrea Costa, Andreu Gual-Grau, Pol Herrero, Helena Torrell, Núria Canela, Lluís Arola

## Abstract

Obesity and its comorbidities are currently considered an epidemic, and the involved pathophysiology is well studied. Recently, the gut microbiota has emerged as a new potential therapeutic target for the treatment of obesity. Diet and antibiotics are known to play crucial roles in changes in the microbiota ecosystem and the disruption of its balance; therefore, the manipulation of gut microbiota may represent a strategy for obesity treatment. Fecal microbiota transplantation, during which fecal microbiota from a healthy donor is transplanted to an obese subject, has aroused interest as an effective approach for the treatment of obesity. To determine its success, a multiomics approach was used that combined metagenomics and metaproteomics to study microbiota composition and function.

To do this, a study was performed in rats that evaluated the effect of a hypercaloric diet on the gut microbiota, and this was combined with antibiotic treatment to deplete the microbiota before fecal microbiota transplantation to verify its effects on gut microbiota-host homeostasis. Our results showed that a high-fat diet induces changes in microbiota biodiversity and alters its function in the host. Moreover, we found that antibiotics depleted the microbiota enough to reduce its bacterial content. Finally, we assessed the use of fecal microbiota transplantation as an obesity therapy, and we found that it reversed the effects of antibiotics and reestablished the microbiota balance, which restored normal functioning and alleviated microbiota disruption.

## Introduction

Obesity is defined as a disequilibrium in energy balance and is currently a global health problem in Western societies, where its prevalence has increased considerably in recent years. Obesity triggers a vast number of comorbidities associated with hypertension, cardiovascular disease, and diabetes, as well as other conditions [1]. It is widely known that obesity is affected by numerous factors, such as diet, lifestyle and genetic background [2], and recently it has been shown to be related to gut microbiota [3], which have been implicated in energy homeostasis and metabolic functions [4]. Moreover, the same factors that affect obesity can modulate gut microbiota composition, and the function of the gut microbiota will be affected by factors involved in gut microbiota-host equilibrium [5].

Several diet-induced animal models of obesity can be used to explore the mechanisms involved in obesity. There are different obesogenic diets that can be employed. One example of these diets is the semi-purified high-fat diet [6,7]. These types of diets are more commonly used in these models due to their well-defined nutritional composition [8–10].

Alterations in the gut microbiota composition have been shown to result in an imbalance that leads to dysbiosis, which likely will have dramatic effects on the maintenance of health [11]. Fecal microbiota transplantation (FMT) is a new and straightforward therapy that manipulates the gut microbiota by transferring healthy donor microbiota into an existing disrupted gut microbial ecosystem. This therapy can be an effective approach to obesity treatment [12,13]. Even though FMT has some limitations, several studies have tested its effectiveness and have demonstrated an improvement of some comorbidities associated not only with obesity [14] but also with other noncommunicable diseases [15–18].

Animal models have been increasingly employed to investigate the role and function of gut microbiota, and there have been several studies where mice that were fed a high-fat diet showed a clear disruption in their microbiota composition [19,20]. Such changes in the microbiota due to diet can modulate important metabolic functions, including fat storage [21]. Among the different animal models available, germ-free (GF) mice represent the model that is most used to study the interaction between hosts and their microbiota, and it is also the preferred option for FMT studies. However, GF mice have less body fat in comparison with wild-type mice, even if they consume more food [22,23], and as a result, they are not the most realistic model for obesity-induced studies. In addition, these animals must be bred in sterile environments, and conducting these kinds of experiments requires skilled personnel and a special infrastructure. Thus, gut microbiota depletion by using a cocktail with a combination of broad-spectrum antibiotics [24] is an accessible alternative to the use of GF mice to study the role of microbiota in the host [18].

Nonetheless, the effect of FMT on hosts has hardly been studied due to its novelty, and specific tools are needed to comprehend these effects. Recently, multiomics approaches have been proposed as the most accurate methods for the study of the complexity of the gut microbiota and its environment [4,21,25]. Metagenomics, which provides a taxonomical profile of the biodiversity present in each experimental condition, and metaproteomics, which is focused on the characterization of the whole proteome to reveal its functionality in the host [5,26,27], are the most promising omics strategies that could be used to reveal the role of gut microbiota.

Hence, the aims of this study were to investigate the role of the gut microbiota with a multiomic approach that combined metagenomics and metaproteomics to determine the effects of a dietary intervention consisting of two different diets (a low-fat diet (LFD) and a high-fat diet (HFD)), to assess the effects of antibiotics on gut microbiota depletion and to corroborate the effectiveness of FMT in rats fed a HFD.

## Materials and methods

### Animals

Forty eight-week-old male Wistar rats (Charles River Laboratories, Massachusetts, USA) were housed individually at 22°C with a light/dark cycle of 12 hours and were given access to food and water *ad libitum* during the experiment. After one week of adaptation, the animals were divided into five groups (n=8). For 9 weeks, two groups were fed a LFD (10% fat, 70% carbohydrate, and 20% protein; D12450K, Research Diets, New Brunswick, USA) or a HFD (45% fat, 35% carbohydrate, and 20% protein; D12451, Research Diets, New Brunswick, USA). Two other groups were also fed either a LFD or a HFD for 11 weeks, and during the last 2 weeks, they were given antibiotic treatment (ABS). The last group was fed a HFD for 14 weeks; at 10 and 11 weeks the rats received antibiotic treatment, and during the last three weeks (12-14), they received FMTs from the LFD group (Fig 1).

**Fig 1.**
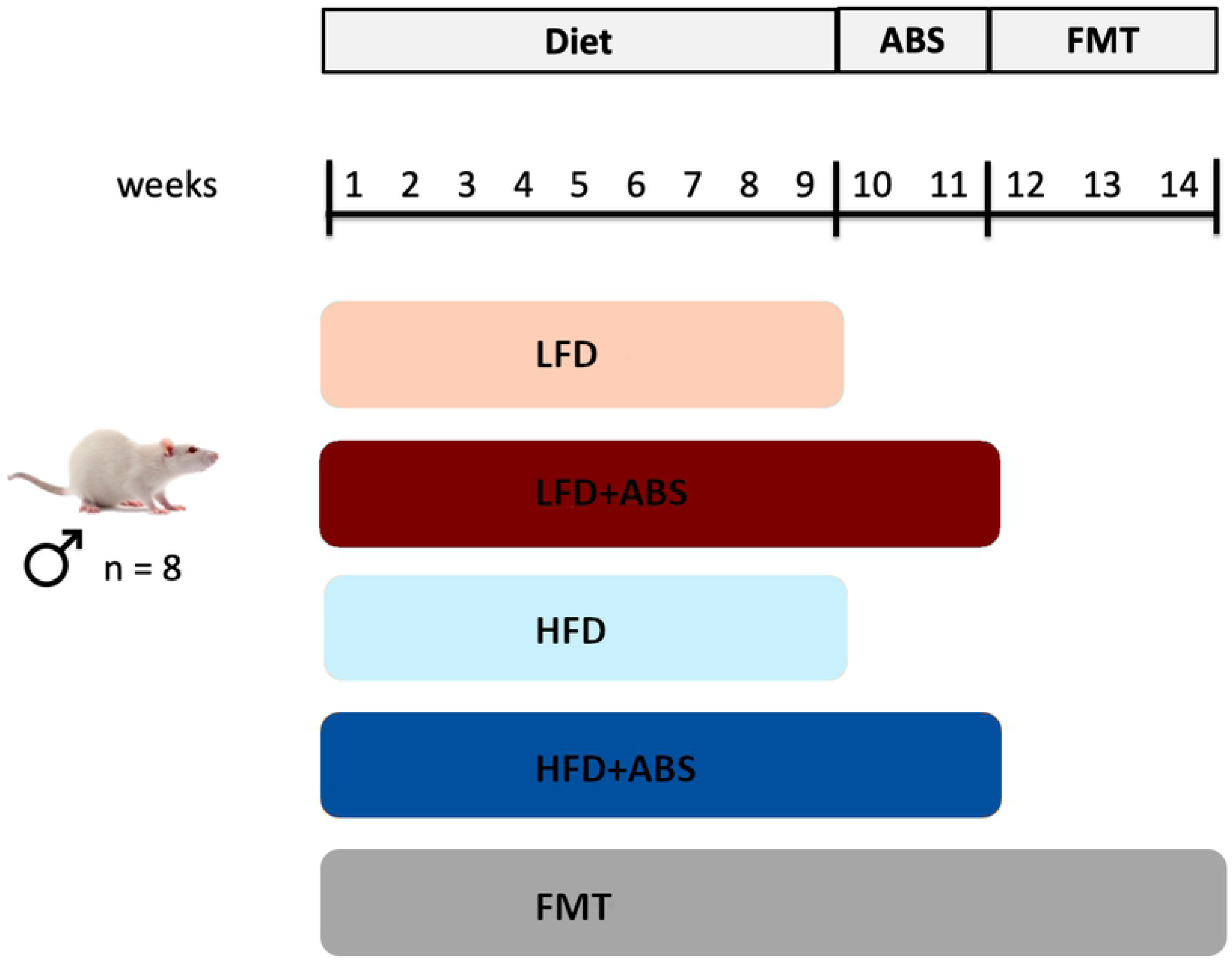
Schematic representation of the experimental design. LFD, Low-Fat Diet; HFD, High-Fat Diet; ABS, Antibiotics; FMT, Fecal Microbiota Transplantation.

Body weight and food intake were measured weekly throughout the study. The body fat mass was determined on weeks 1, 9, 11 and 14 by nuclear magnetic resonance (NMR) using an EchoMRI-700^TM^ device (Echo Medical Systems, L.L.C., Houston, USA).

Cecal samples were obtained immediately after the animals were sacrificed and were frozen in liquid nitrogen and stored at −80°C until the analyses were performed.

The Animal Ethics Committee of the University of Rovira i Virgili (Tarragona, Spain) approved all procedures.

### Antibiotic treatment and Fecal Microbiota Transplants

ABS was started 9 weeks after being fed either diet, and the cocktail of antibiotics used consisted of 0.5 g/L vancomycin (Sigma-Aldrich, UK) and 1 g/L neomycin, metronidazole and ampicillin (Sigma-Aldrich, UK). The water flasks were supplemented with the antibiotic cocktail. The mixture was freshly prepared every day, and the animals were given free access to it.

The FMTs were performed for 3 weeks; cecal content from LFD-fed rats was administered by oral gavage for four consecutive days during the first week, two alternating days during week 2 and three days before the rats were sacrificed. Omeprazole (20 mg/kg) was administered by oral gavage 4 hours before each FMT.

### Metagenomics analysis

#### DNA extraction and 16S rRNA gene amplification and purification

DNA was extracted from 300 mg cecal samples using a QIAamp DNA Stool Mini Kit (Qiagen Inc., Hilden, Germany) according to the manufacturer’s instructions. The DNA purity and integrity were assessed using spectrophotometry (NanoDrop, Thermo Fisher Scientific, Massachusetts, USA).

Two variable regions (V3 and V4) in the 16S rRNA gene were amplified by PCR as described previously [21].

#### Ion Torrent sequencing and taxonomic assignments

A multiplexed mixture of twenty DNA samples was diluted to a concentration of 60 pM prior to clonal amplification. The Ion 520 & Ion 530 Kit-Chef (Life Technologies, California, USA) was employed for template preparation and sequencing according to the manufacturer’s instructions. The prepared samples were loaded on an Ion 530 chip and sequenced using the Ion S5 system (Life Technologies, California, USA).

After sequencing, the Ion Torrent Suite software package was used to remove the low-quality and polyclonal sequences, and the remaining reads were then analyzed using QIIME [28]. The analysis included OTU (operational taxonomic unit) clustering, alpha diversity analysis, OTU analysis and species annotation (OTU table), and beta diversity analysis. The OTU table, which indicates the number of reads per sample per OTU, was used for the subsequent statistical analysis.

### Metaproteomics analysis

The metaproteomics methodology was conducted as described previously [21,29] with minor modifications.

#### Cell lysis and protein digestion

Briefly, 300 mg of stool sample was subjected to differential centrifugation to collect the microbial cells, and the obtained bacterial pellet was suspended in SDS-extraction buffer (2% SDS, 100 mM DTT, and 20 mM Tris-HCl pH 8.8), incubated at 95°C and subjected to a bead beating process (Bullet Blender, Cultek, Barcelona, Spain) combined with freeze-thawing cycles. The proteins were purified by TCA/acetone precipitation, and 75 μg of protein from each sample was reduced and alkylated, loaded onto a polyacrylamide gel and digested overnight at 37°C with trypsin (Promega, Wisconsin, USA) at an enzyme-to-protein ratio of 1:100.

#### Peptide TMT 10plex labeling

The digested proteins were desalted with an HLB SPE column before labeling with TMT 10plex reagent (Thermo Fisher Scientific, Massachusetts, USA) according to the manufacturer’s instructions. To normalize the samples and the TMT batches, all samples were pooled and labeled with a 126-tag, and the pooled sample was included in each batch. Then, the labeled peptides from each sample were mixed together and desalted again with an HLB SPE column.

#### Peptide fractionation

The pooled samples were fractionated by isoelectric focusing with an Off-Gel Fractionator (OG) (Agilent Technologies, California, USA) and 24-well IPG strips (with a nonlinear gradient from pH 3 to pH 10) according to the manufacturer’s protocol. After fractionation, each of the 24 fractions was desalted again with an HLB column (Waters, Massachusetts, USA) prior to nanoLC-Orbitrap MS/MS analysis.

#### nanoLC-Orbitrap MS/MS analysis

The 72 fractions obtained from the OG fractionation (3 TMT x 24 fractions) were loaded on a trap nanocolumn (0.01 x 2 cm, 5 μm; Thermo Fisher Scientific, Massachusetts, USA) and separated with a C-18 reversed-phase (RP) nanocolumn (0.0075 x 12 cm 3 μm; Nikkyo Technos Co. LTD, Japan). The chromatographic separation was performed with a 90-min gradient that used Milli-Q water (0.1% FA) and ACN (0.1% FA) as the mobile phase at a rate of 300 nl/min. Mass spectrometry analyses were performed on a LTQ-Orbitrap Velos Pro (Thermo Fisher Scientific, Massachusetts, USA) by acquiring an enhanced FT-resolution spectrum (R=30,000 FHMW) followed by the data-dependent FT-MS/MS scan events (FT-(HCD)MS/MS (R=15,000 FHMW at 35% NCE) from the ten most intense parent ions with a charge state rejection of one and a dynamic exclusion of 0.5 min.

The 24 raw data files for each TMT-plex were analyzed by multidimensional protein identification technology (MudPIT) using Proteome Discoverer software v.1.4.0.288 (Thermo Fisher Scientific, Massachusetts, USA). For protein identification, all MS and MS/MS spectra were analyzed using the Mascot search engine (version 2.5), which was set up to search two different SwissProt databases based on (i) *Rattus norvegicus* (8,003 sequences) and (ii) an in-house metagenomics database created from metagenomics results at the family level using the Uniref100 sequence identity to reduce the database size and avoid false positive findings (23,768,352 sequences). Two missed cleavage sites were allowed by assuming trypsin was used for digestion, and an error of 0.02 Da for the FT-MS/MS fragment ion mass and of 10.0 ppm for the FT-MS parent ion mass was allowed. For TMT-10plex analysis, lysine and the N-termini were set as quantification modifications, while methionine oxidation and the acetylation of N-termini were set as dynamic modifications, and carbamidomethylation of cysteine was set as a static modification. The false discovery rate (FDR) and protein probabilities were calculated by a fixed PSM validator.

For protein quantification, the ratios between each TMT label and the 126-TMT label were used and normalized based on the protein median.

### Statistical Analysis

To determine the significant metagenomic and protein changes between the different conditions under study, the Mass Profiler Professional Software v.14.5 (Agilent Technologies, Massachusetts, USA) was used. The data were log2 transformed and mean-centered for the multivariate analysis (Principal Component Analysis, PCA) and the univariate statistical analysis (Student’s t-test).

## Results and discussion

### Effect of diet, ABS and FMT on body weight and fat mass

During the nine-week period, the HFD group exhibited a significant increase in body weight after week 5 (p<0.001), and the body fat mass measured by NMR in this group was also significantly higher. During antibiotic administration (from 10 to 11 weeks), both parameters were decreased but remained significantly different between the different dietary groups (p < 0.001). Nonetheless, when the FMTs began, the animals recovered their body weight, and their body mass content increased again (Fig 2). Similar results have been reported in other studies in which rats were fed a commercial diet with a high fat content [8,30]; moreover, when an antibiotic treatment is administered, normally the animals exhibit reductions in appetite and food intake, and a loss of body weight occurs [31–33]. This could be an explanation for the fact that the rats lost body weight during antibiotic treatment.

**Fig 2.**
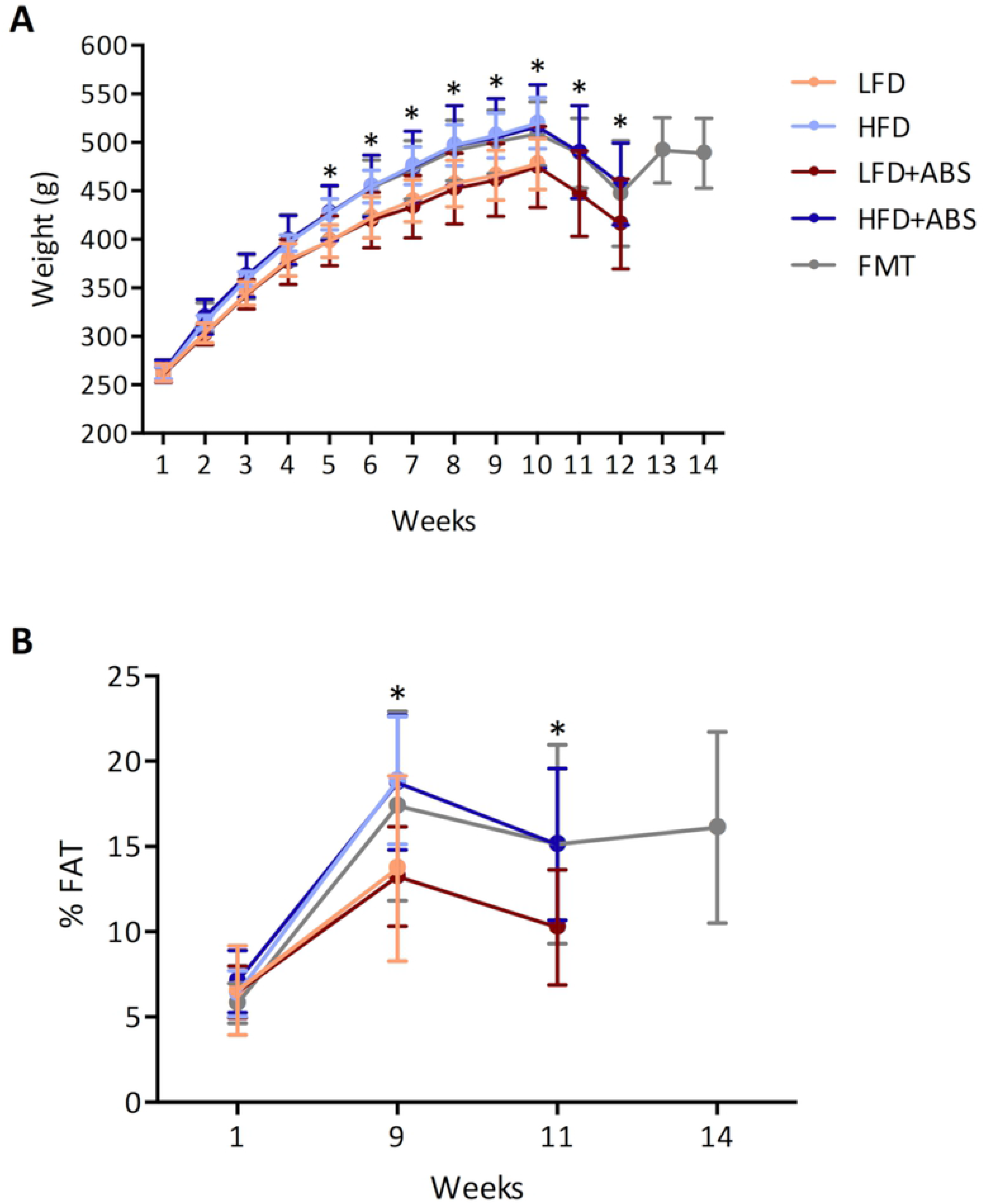
A) Measurement of body weight during the 14 weeks of study and B) the percentage of body fat measured at weeks 1, 9, 11 and 14. *p<0.05, calculated using ANOVA.

### Changes in microbiota biodiversity and functionality caused by diet

To assess the impact of diet on gut microbiota, metagenomics and metaproteomics analyses were performed in cecal samples obtained immediately after sacrifice that were divided into two equal portions for use in both omics approaches.

The metagenomics analysis was performed using NGS Ion Torrent Technology. The sequencing runs produced a total of 49,106,850 reads that were filtered for quality, and 25,535,562 were obtained for the QIIME analysis. From these reads, a total of 17,220 OTUs were obtained for the V3 and V4 regions of the 16S rRNA gene sequence that were used to analyze the relative abundances and diversity in the microbiota at different taxonomical levels.

The two dominant phyla were Bacteroidetes (14.6 – 44.5%) and Firmicutes (52.6 – 84.1%) in both groups, which is in line with the results of previous studies [21,34]. The Bacteroidetes to Firmicutes ratio (B/F) was significantly increased (p=0.023) in the HFD group (B/F=0.393) compared to the LFD group (B/F=0.294). No differences were found with regard to phylum level or alpha diversity or the Shannon and Simpson indexes, but some comparisons could be made at the family level, and a clear separation between the groups was observed (Fig 3). In total, the abundances of 9 family taxa were significantly different between the LFD- and HFD-fed rats (Table 1); 7 out of 9 were from the Firmicutes phylum, and 6 were from the Clostridiales order. Even the abundance of Clostridiales was not very high (52.0 – 89.8%), and Clostridiales was probably the order most affected by diet and was responsible for gut microbiota disruption. Similar results have been reported before that have shown clear differences in microbiota composition when a diet rich in fat is administered [19–21,35] and that the Firmicutes phylum, including the Clostridiales order, represents the taxon with the most changes in terms of microbiota composition.

**Fig 3.**
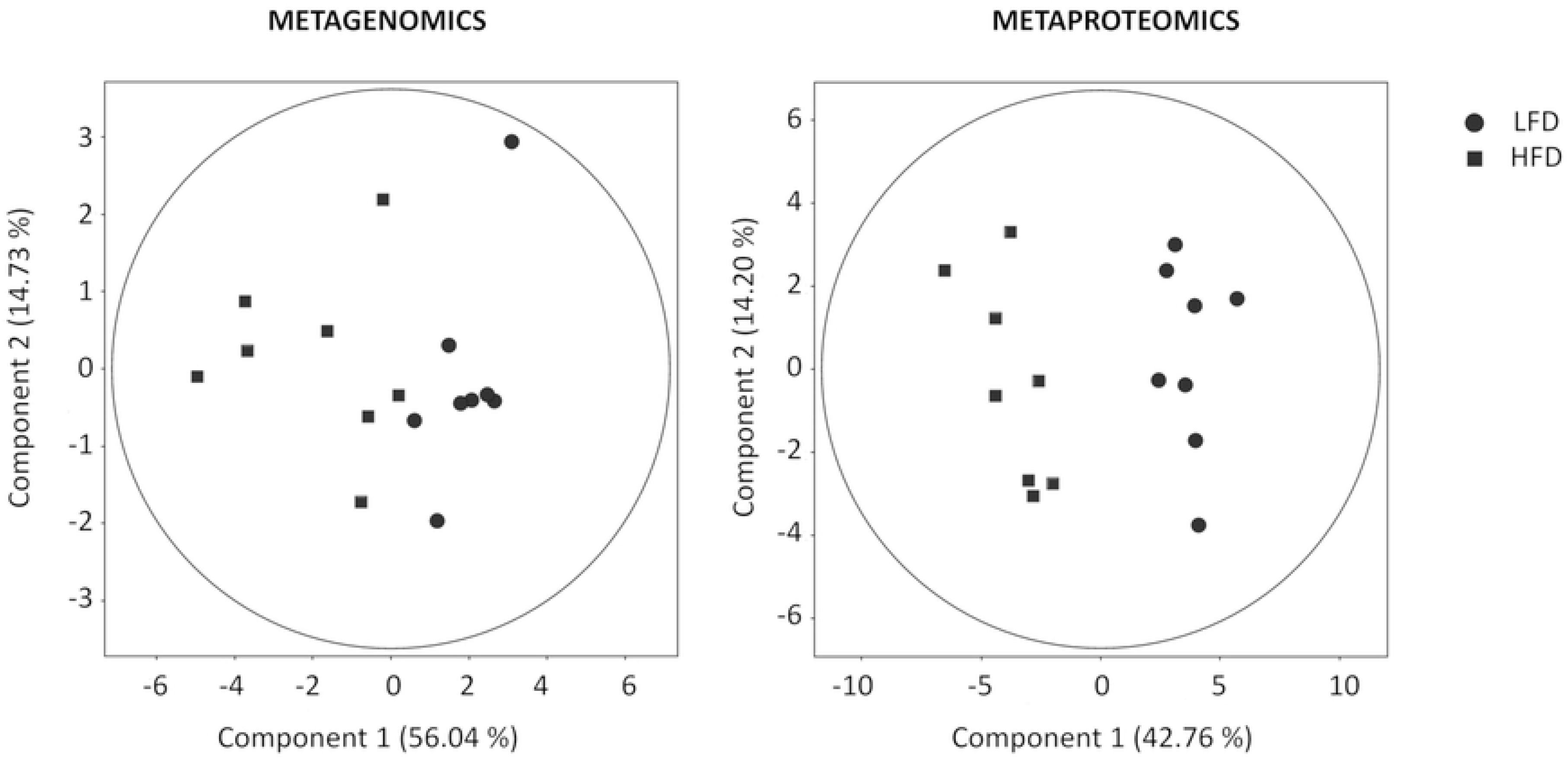
A) PCA of the family OTU abundance and the proteins that were identified that shows the separation between the LFD and HFD groups. The first two components are shown along with the percent variance that is explained by each. The points correspond to the individual samples.

**Table 1:** Families with significant differences in abundance between the LFD and HFD groups. Metagenoics analysis.

| Family | Phylum | LFD (%) | HFD (%) | Regulation | FC | p-value |
| --- | --- | --- | --- | --- | --- | --- |
| Coriobacteriaceae | Actinobacteria | 0.011 | 0.007 | down | -2.44 | 0.029 |
| Streptococcaceae | Firmicutes | 0.026 | 0.012 | down | -2.67 | 0.005 |
| Christensenellaceae | Firmicutes | 0.196 | 0.116 | down | -1.86 | 0.004 |
| Clostridiaceae | Firmicutes | 0.268 | 0.088 | down | -5.07 | 0.008 |
| Dehalobacteriaceae | Firmicutes | 0.017 | 0.142 | up | 5.48 | 0.017 |
| Peptostreptococcaceae | Firmicutes | 0.188 | 0.109 | down | -6.47 | 0.007 |
| Veillonellaceae | Firmicutes | 2.750 | 1.706 | down | -10.86 | 0.028 |
| Mogibacteriaceae | Firmicutes | 0.046 | 0.019 | down | -2.46 | 0.001 |
| Desulfovibrionaceae | Proteobacteria | 0.810 | 0.462 | down | -1.66 | 0.039 |

In addition to metagenomics, metaproteomics was performed to assess the impact of diet on microbiota function. A total of 72 fractions were analyzed, and 1598 bacterial proteins were identified. By filtering these proteins on the basis of their being present in at least 50% of samples from at least one of the groups, 415 were selected due to greater confidence.

To assess the impact of diet, the differences between the LFD and HFD groups were identified, which showed that 22 and 11 proteins were up-regulated in the HFD group and the LFD group, respectively (Table 2); most of these proteins were involved in metabolic functioning and played roles in numerous biological processes, such as the TCA and ATP metabolic pathways. The differences in all 33 of these proteins allowed us to perfectly separate both groups (Fig 3).

**Table 2.**
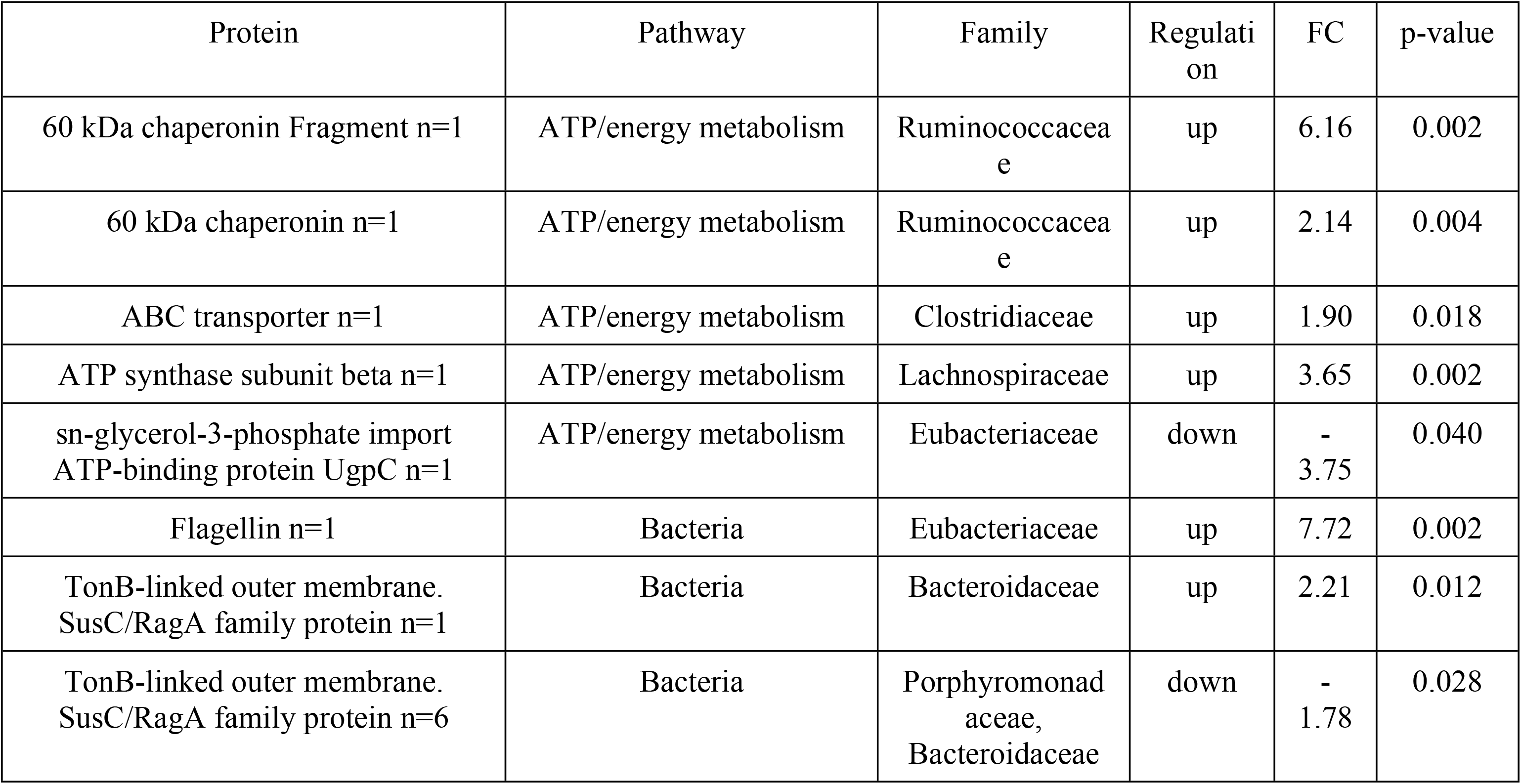
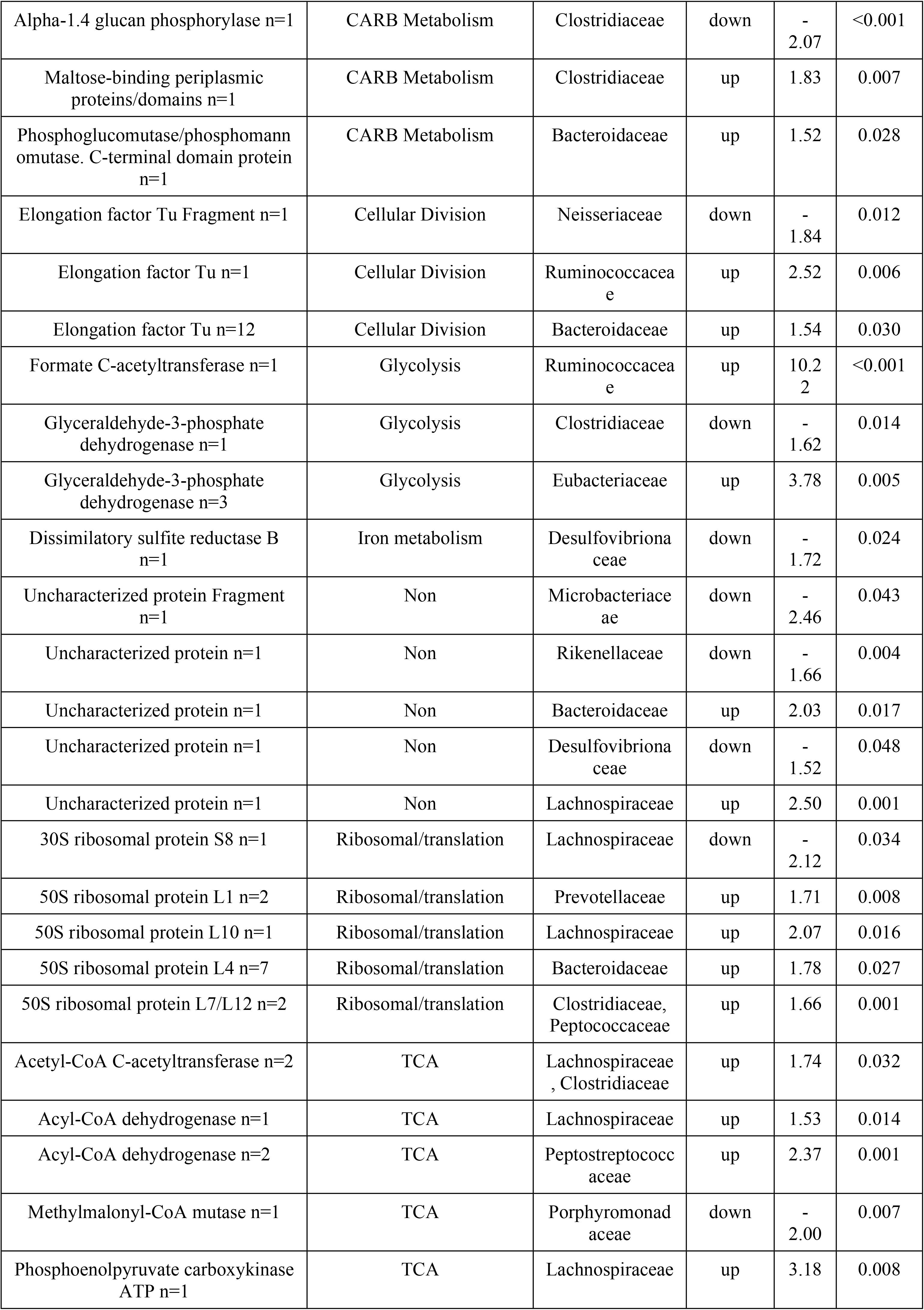
Proteins significantly up- or downregulated between the LFD and HFD groups. Metaproteomics analysis.

Moreover, each protein was assigned to a family taxon, and if we compare the phyla and orders corresponding to the significant proteins we found, we find that all such proteins were derived from Firmicutes and Clostridiales, which is in line with the metagenomics analysis. However, the correlation between the metagenomics and metaproteomics analyses was not complete at the family taxa level.

Few studies have used similar experimental approaches that can corroborate our results. Only a single previous study carried out in our laboratory assessed the functionality of microbiota [21], and other studies performed in mice have explored the changes in the main functions of the microbiota resulting from a high-fat diet [19]. In both cases, the majority of the functions affected by microbiota were involved in important metabolic functions.

### The microbiota composition was disrupted by antibiotic treatment

After the assessment of diet, a metagenomics analysis was performed in each dietary group after ABS treatment to verify microbiota disruption, and the results were compared to those from groups without ABS administration. The LFD group was compared to the LFD+ABS group, and the HFD group was compared to the HFD+ABS group.

The actual OTU abundance is shown in Fig 4A, which shows that the majority of the bacterial content in the cecal microbiota was depleted by antibiotic treatment and also indicates that the bacterial content was restored by FMT.

**Fig 4.**
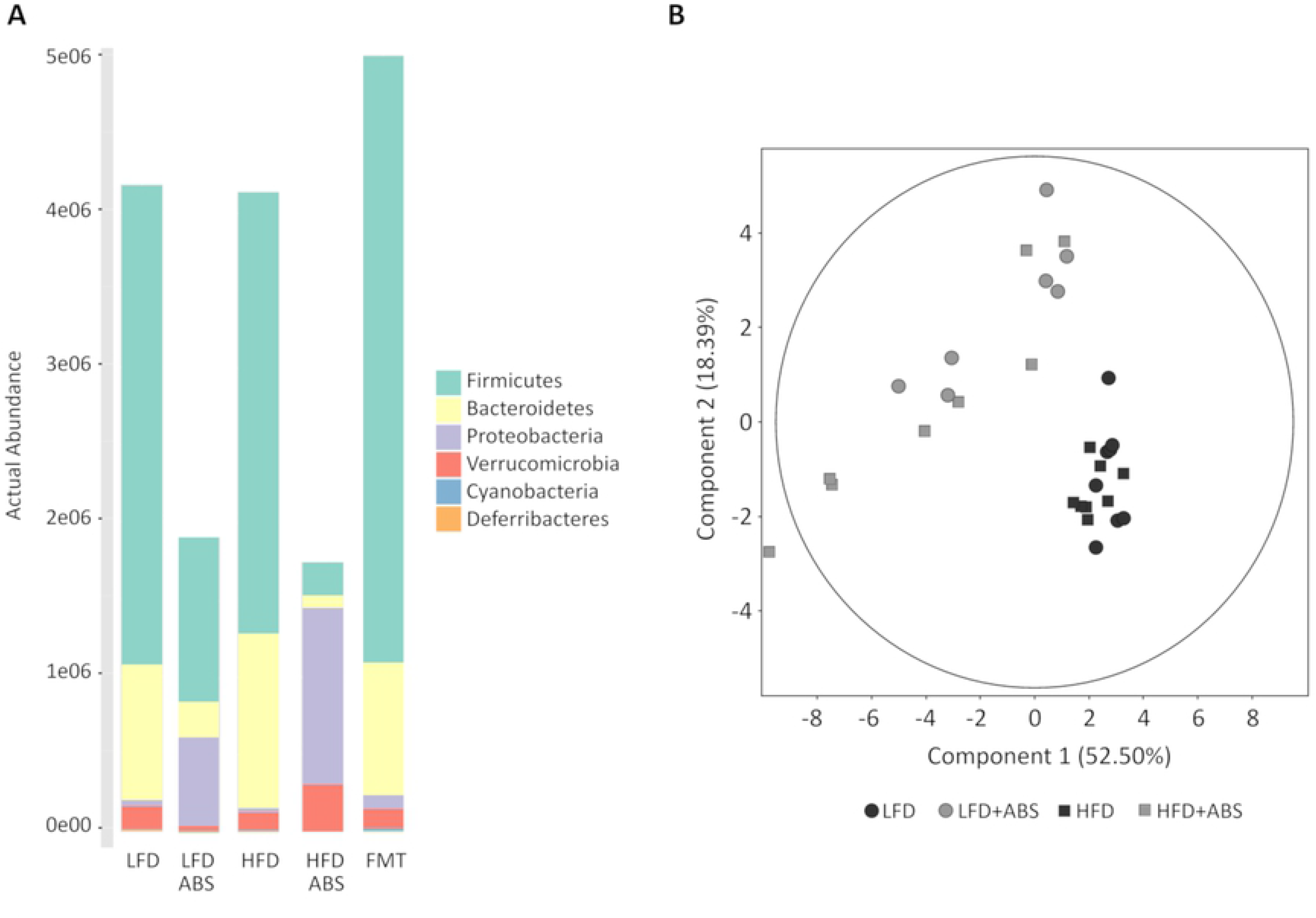
A) Differences in the actual OTU abundances among the different groups at the phylum taxon levels. B) PCA of the differences between the groups treated with and without ABS. The first two components are shown along with the percentages of variance that they explain. The points correspond to individual samples.

Regarding the metagenomics results in the LFD groups, a decrease in both Bacteroidetes (LFD 21.2%; LFD+ABS 10.5%) and Firmicutes (LFD 74.0%; LFD+ABS 45.9%) phyla were observed in the LFD+ABS group, whereas Proteobacteria (LFD 1.0%; LFD+ABS 38.9%; p=0.007) represented the dominant phylum after antibiotic treatment. The B/F ratio was significantly decreased (LFD B/F=0.294; LFD+ABS B/F=0.216; p=0.039); furthermore, the alpha diversity based on observed OTUs and the Shannon and Simpson indexes was not significant (p=0.192, p=0.095, and p=9.982, respectively).

In contrast to the LFD-fed rats, in the HFD-fed rats no significant differences were found for the B/F ratio (HFD B/F=0.393; HFD+ABS B/F=0.382; p=0.110), although similar results due to antibiotic treatment occurred in the LFD+ABS group, which resulted in dramatic decreases in both phyla (Bacteroidetes, HFD 26.4%, HFD+ABS 8.70%; Firmicutes, HFD 70.5%, HFD+ABS 26.5%) and an enormous increase in Proteobacteria (HFD 0.6%; HFD+ABS 59.1%; p<0.001). Moreover, the alpha diversity decreased significantly in the HFD+ABS group compared to the HFD group (observed OTUs p=0.011; Shannon index p=0.009; Simpson index p=0.011).

The results found in both ABS groups were consistent with those of previous studies where a similar cocktail of ABS was administered to wild-type mice, which caused the relative abundances of Bacteroidetes and Firmicutes to decrease, while the abundances of Proteobacteria and Cyanobacteria increased [24,33]. These results could be explained by differences in antibiotic effectiveness against bacteria from different phyla. As can be seen, the antibiotics had stronger activity against bacteria from the Bacteroidetes and Firmicutes phyla and are the reason why changes in the abundance of Proteobacteria were observed in these particular groups, although the bacterial DNA quantity was considerably lower in the ABS samples.

In addition, the relative abundances were compared at the family level for both the LFD and HFD groups. In the LFD group comparison, distinct differences were found in 18 family taxa, most of which are in the Firmicutes and Proteobacteria phyla, whereas in the HFD groups distinct differences were found for 15 family taxa among the Bacteroidetes, Firmicutes and Proteobacteria phyla (Table 3 and Fig 4B). A total of 10 families were found to be in common in rats fed both diets: *S24-7* from Bacteroidetes; *Enterococcaceae*, *Streptococcaceae*, *Christensenellaceae, Lachnospiraceae, Peptococcaceae* and *Ruminococcaceae* from Firmicutes; *Alcaligenaceae*, *Enterobacteriaceae* and *Pseudomonadaceae* from Proteobacteria. All of these were regulated equally in the ABS groups compared to the respective non-ABS groups. Similar changes in microbiota composition were found in several previous studies of chronic antibiotic exposure [36–38] that showed a significant reduction of bacterial richness and diversity and corroborated the effectiveness of gut microbiota depletion for FMT studies.

**Table 3:**
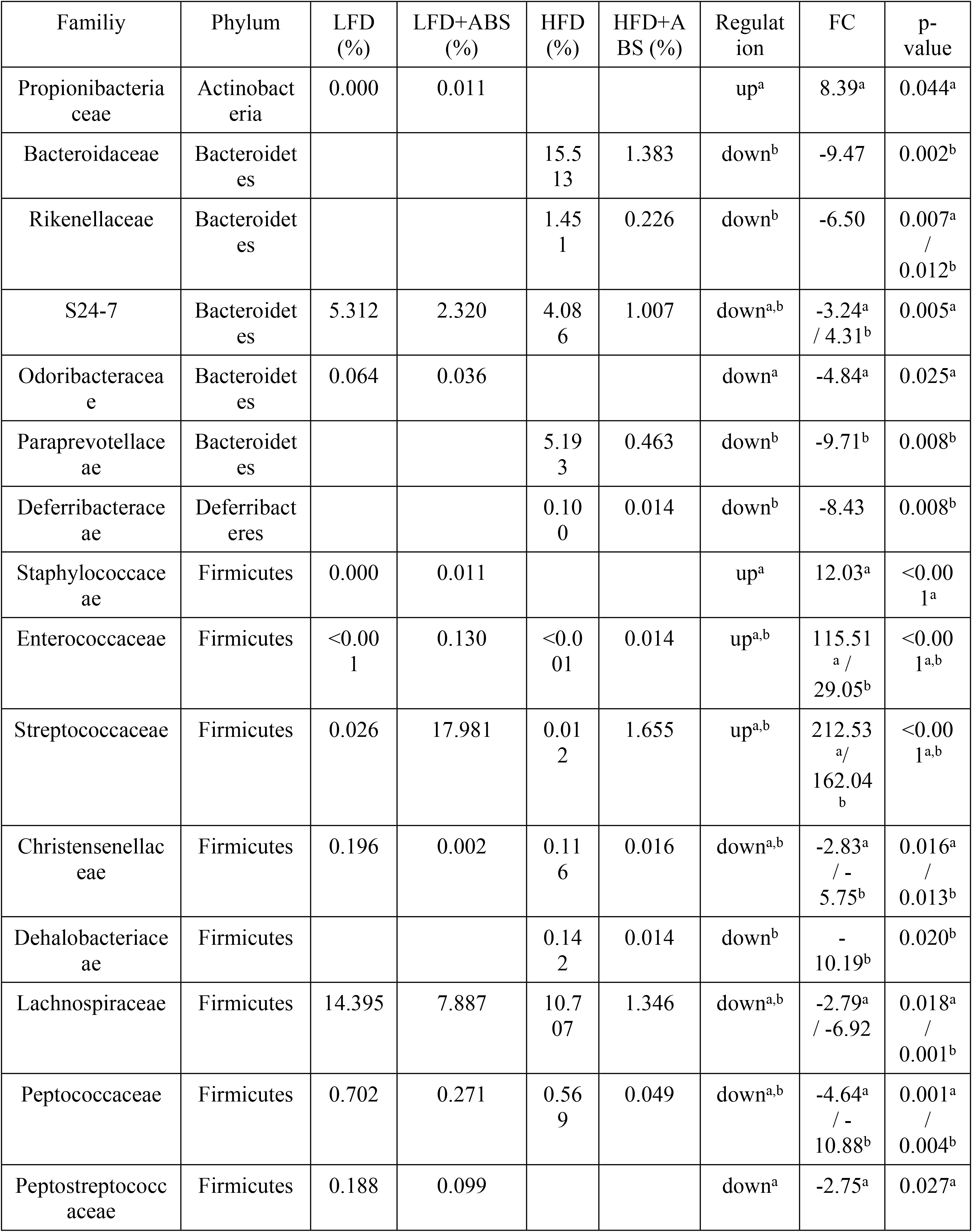
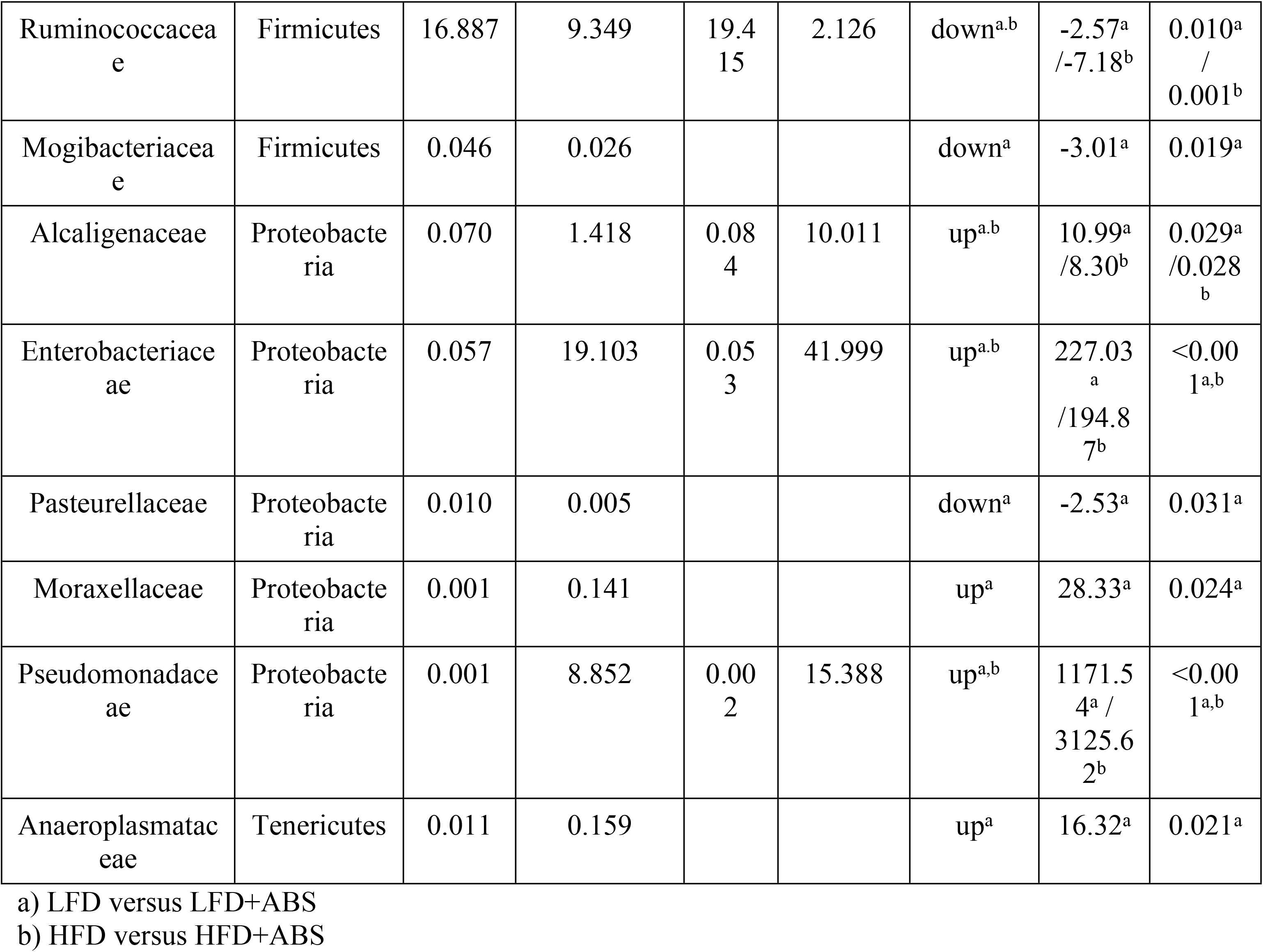
Families with significant differences in abundance between the diet-only groups and the respective diet-plus-ABS groups. Metagenomics analysis

### FMT restored microbiota biodiversity and functionality

To assess the use of FMT as a possible treatment for overweight or obesity caused by dietary habits, FMT was performed by transplanting cecal microbiota content from LFD rats to HFD rats previously depleted by ABS treatment. The metagenomics analysis revealed that Bacteroidetes (LFD 21.2%, HFD 26.4%, FMT 18.0%) and Firmicutes bacteria (LFD 74.0%, HFD 70.5%, FMT 77.0%) were restored after FMT to levels similar to those found prior to antibiotic treatment, and consequently Proteobacteria (LFD 1.0%, HFD 0.6%, FMT 2.0%) were decreased considerably. As can be observed, the B/F ratio was more similar between the LFD (B/F=0.294) and FMT (B/F=0.233) groups since no significant differences were observed (p=0.109), whereas the B/F ratio was significantly different in the HFD group (B/F=0.393, p=0.011). Regarding alpha diversity, some differences were found in terms of the observed OTUs and the Shannon and Simpson indexes (p=0.001, p<0.001, and p<0.001, respectively) in the FMT groups compared to both the LFD and the HFD group (one-way ANOVA). These differences could be ameliorated by prolonging the FMT treatment in future studies, but these differences were quite small.

Additionally, a decision was made to determine whether some differences could be found at the family level. After one-way ANOVA, the abundances of 10 families were found to be significantly different between two of the three experimental groups, as shown in Table 4. However, Tukey post hoc tests revealed that the abundances of only 3 families were significantly different between the LFD and HFD groups (*Clostridiaceae, Christensenellaceae* and *Mogibacteriaceae)* and were not significantly different when the FMT group was compared to the LFD group. Moreover, these three families were equally regulated in the LFD and FMT groups when compared to the HFD group. This indicates greater similarity between the FMT and LFD groups than between either of these groups and the HFD group. PCA and hierarchical cluster analysis (HCA) based on the relative abundances of these 3 families was performed to assess the similarities between the LFD, HFD and FMT groups. As shown in Fig 5A and B, the FMT group was more similar to the LFD group than the HFD group even when the rats were fed a hypercaloric diet during and after ABS and FMT treatment.

**Fig 5.**
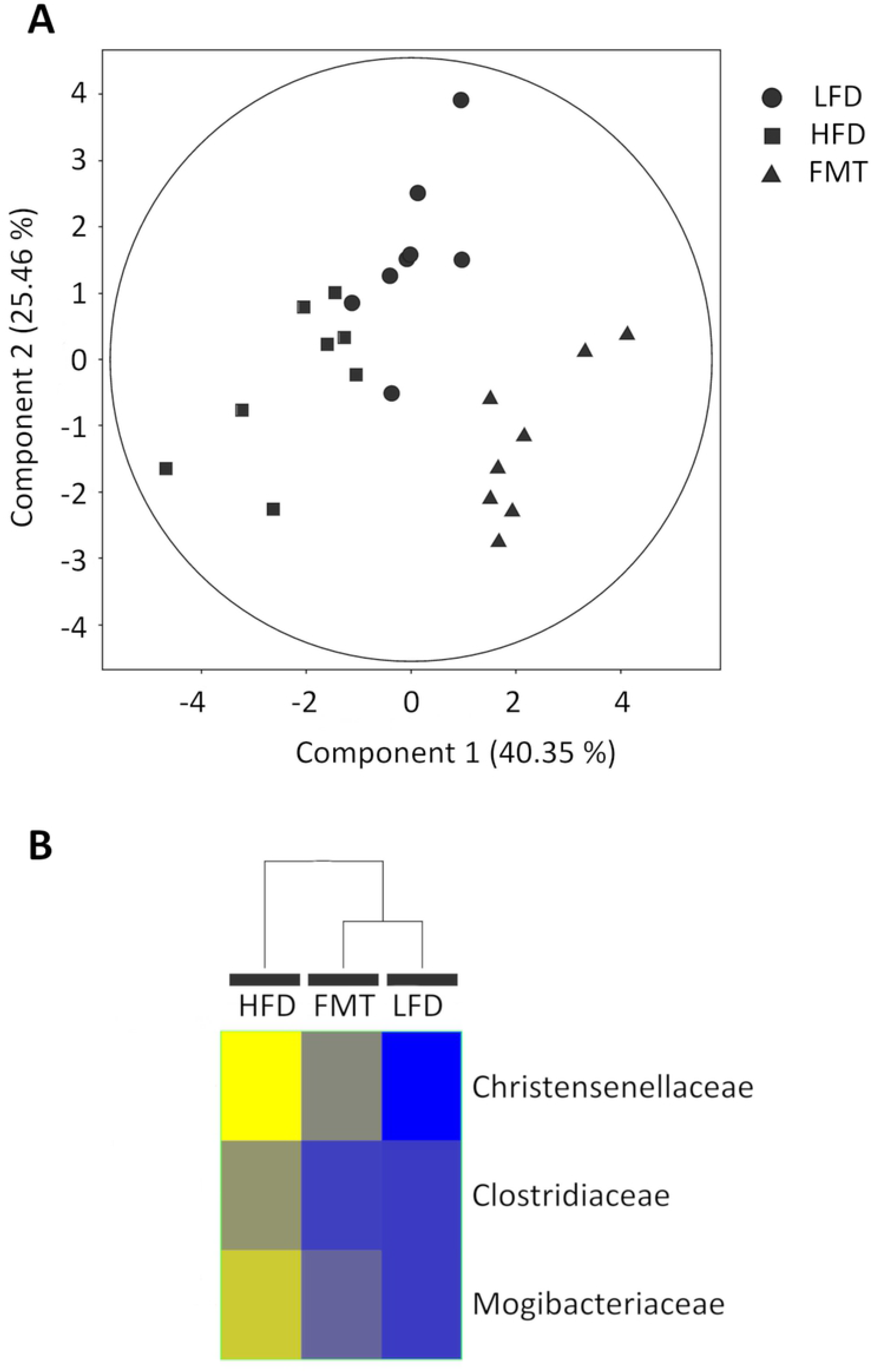
A) PCA of OTU abundance. The first two components are shown along with the percentage of variance that they explain. The points correspond to individual samples. B) Hierarchical clustering analysis of the three significant families in the LFD, HFD and FMT groups.

**Table 4.**
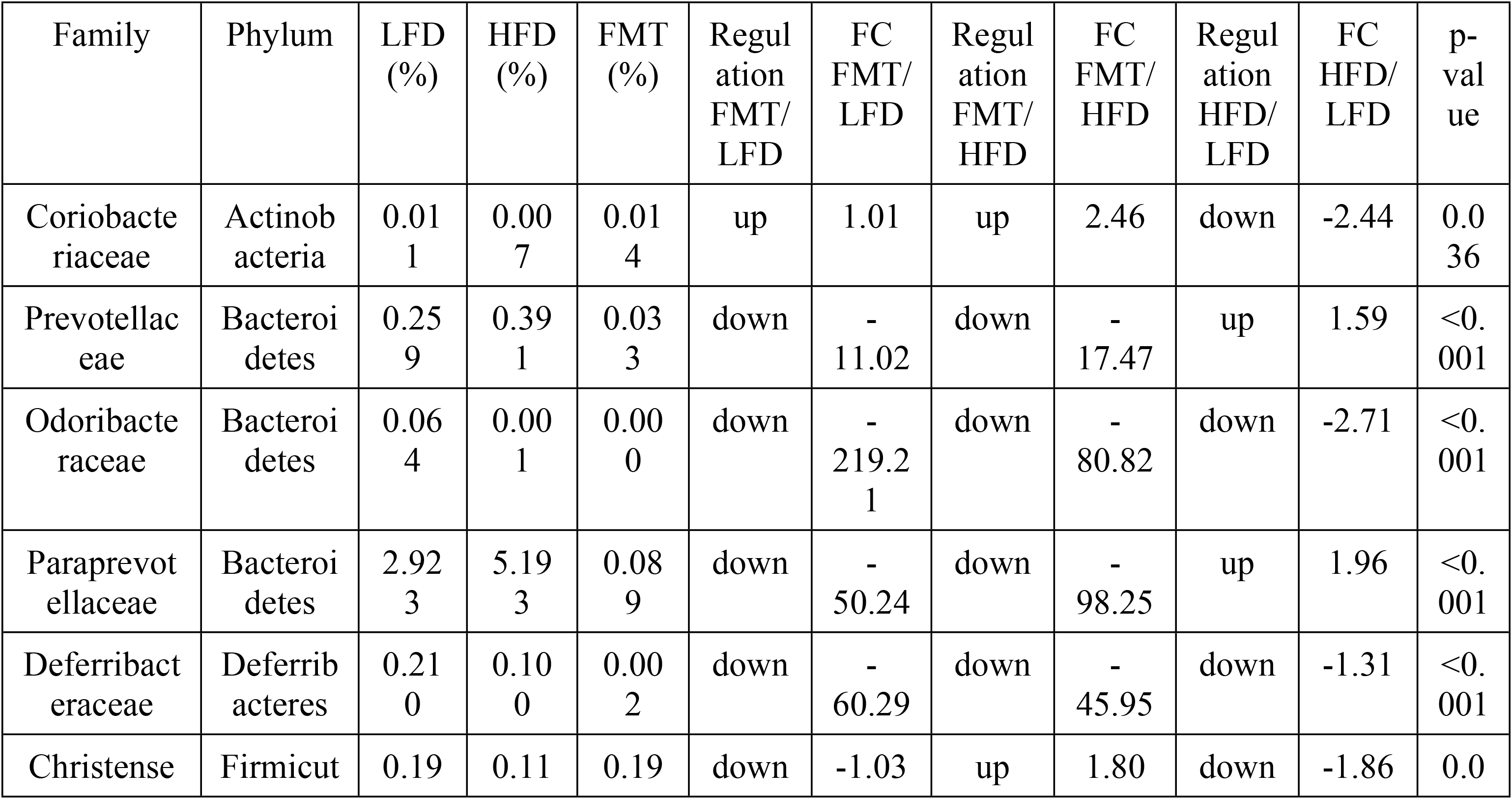
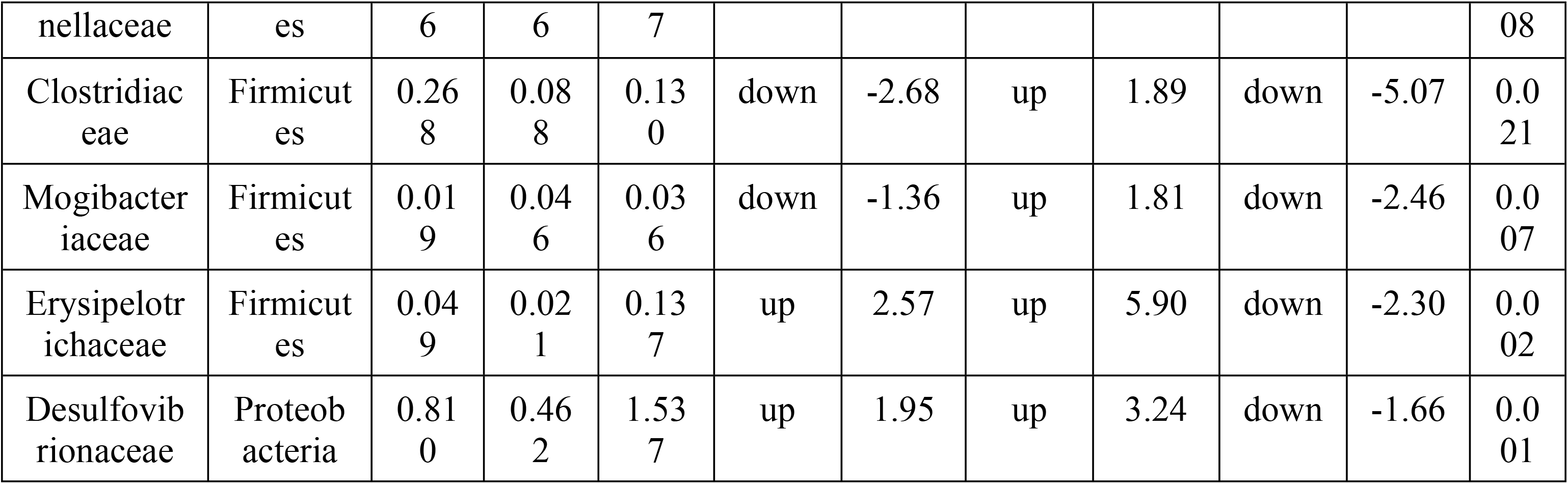
Families with significant differences in abundance based on ANOVA between the LFD, HFD and FMT groups. Metagenomics analysis.

Furthermore, metaproteomics was performed to assess the disruption of microbiota functionality due to diet and FMT. To assess whether there were some differences between these three groups, one-way ANOVA was performed, and a total of 235 proteins were found to be different in one of the groups. These proteins allowed us to perfectly separate these three groups during PCA (Fig 6A). Of these 235 proteins, 155 and 194 were different in the FMT group compared to the LFD and HFD groups, respectively. Moreover, 21 proteins were significantly different between the LFD and HFD groups, and this could also explain the differences between these groups and the FMT group. As seen, the greater number of significant proteins in the FMT group compared to the LFD and HFD groups may be due to the low level of biodiversity observed in the FMT group, as previously described.

**Fig 6.**
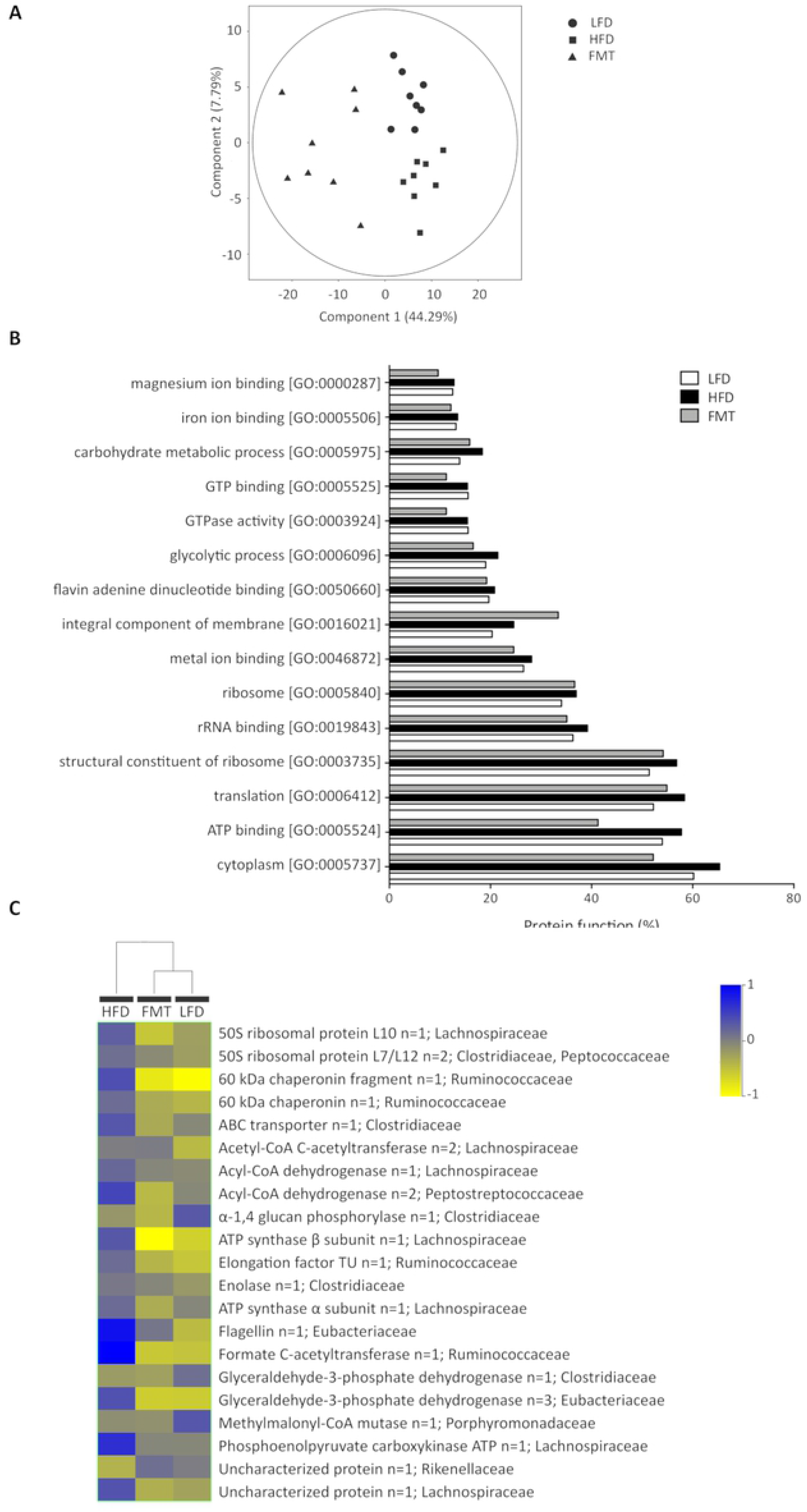
A) PCA of up-and downregulated proteins showing the separation between the three groups. The first two components are shown along with the percentages of variance that they explain. The points correspond to individual samples. B) Percentages of proteins that represent the 15 most abundant protein functions according to Gene Ontology (GO) terms in the LFD, HFD and FMT groups. C) Hierarchical clustering analysis of 21 significant proteins in the LFD, HFD and FMT groups.

Subsequently, each of these proteins was assigned to a taxonomical family to correlate them with the metagenomics results, but only the Clostridiaceae family showed significant differences based on both omics approaches (Table 5). However, if we made the same comparison based on the order taxon, the majority of families that showed significant differences were from the Clostridiales order and the Firmicutes phylum, as was observed when only the LFD and HFD groups were compared.

**Table 5.**
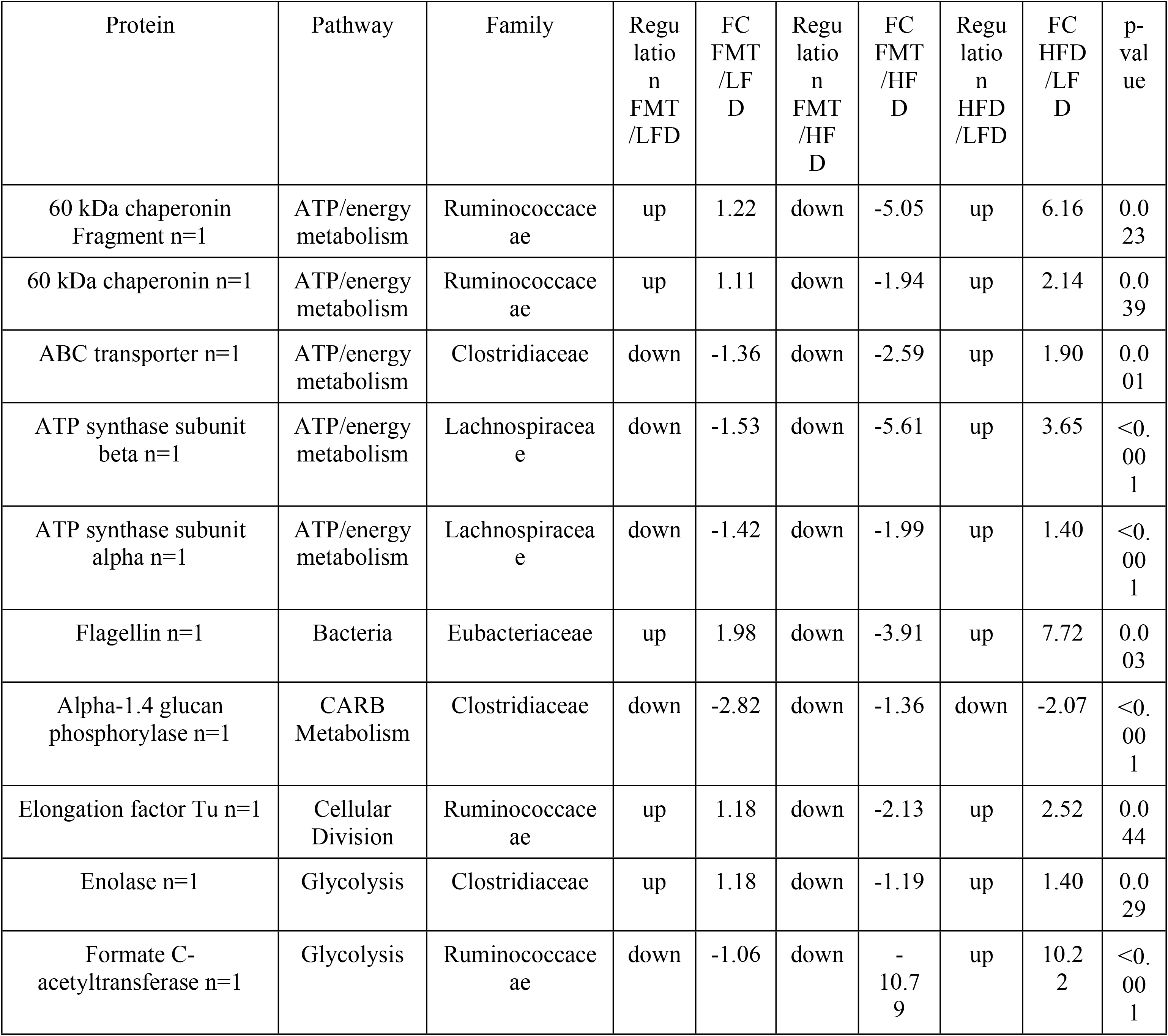
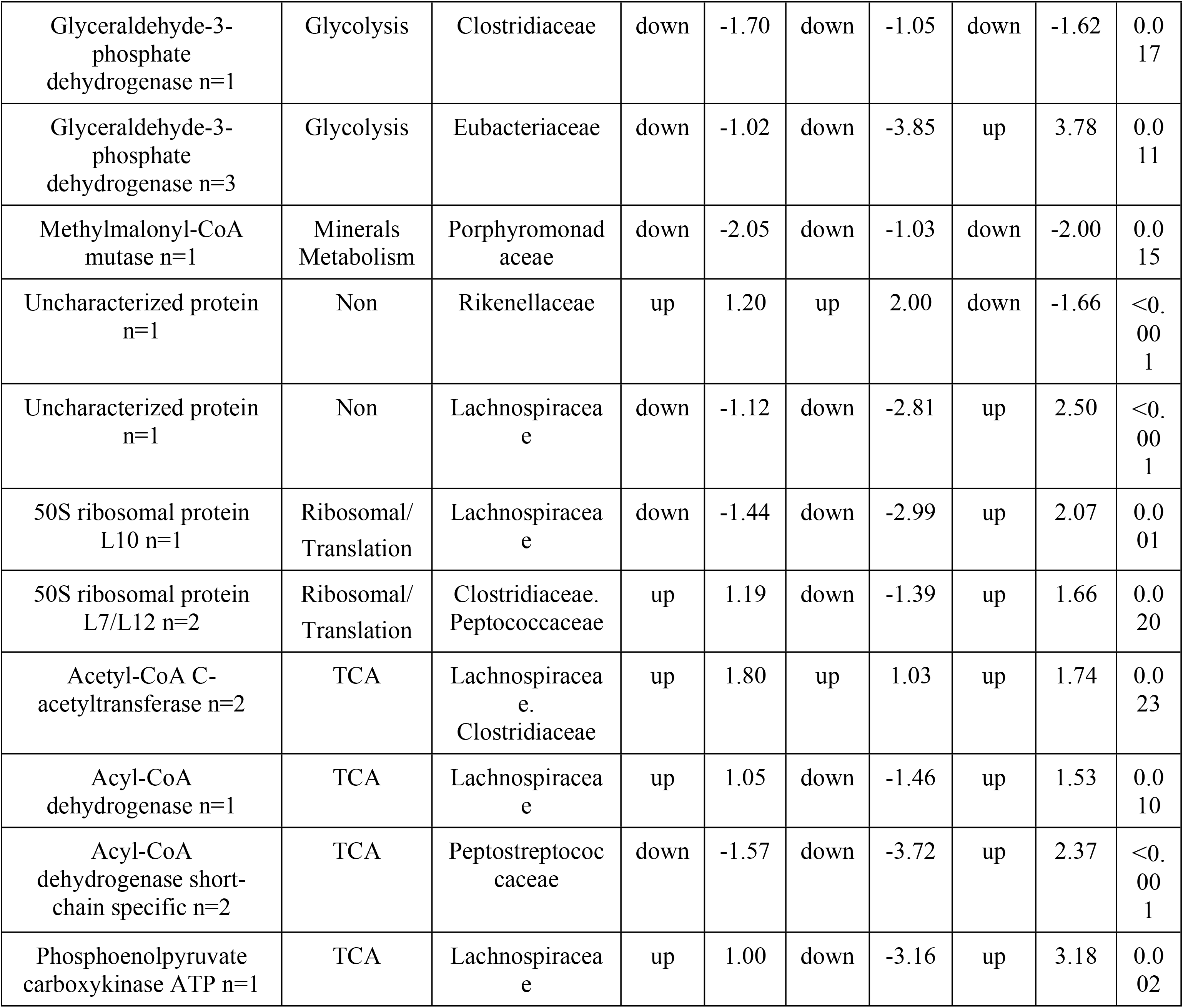
Proteins significantly up- or downregulated between the LFD, HFD and FMT groups. Metaproteomics analysis.

To more deeply understand the function of the whole gut microbiome, GO functions were attributed to each protein, and the 15 most highly represented activities are shown in Fig 6B. The most highly represented functions in each group were represented by proteins involved in the structure of the ribosome [GO:0003735], translation [GO:0006412], the cytoplasm [GO:0005737] and ATP binding [GO:0005524]. These four GO functions were slightly increased in the HFD group, whereas ATP binding and cytoplasmic proteins were decreased in the FMT group compared to both the HFD and LFD groups. Additionally, proteins representing integral components of the membrane [GO:0016021] were increased in the FMT group compared to the other groups.

Based on their metabolic functions, the majority of these proteins were involved in important metabolic pathways such as those involved in ATP/energy metabolism or glycolysis. A total of 8 proteins were equally regulated in both the LFD and FMT groups compared to the HFD group, and important proteins such as Acyl-CoA dehydrogenase and phosphoenolpyruvate carboxykinase, both from the Lachnospiraceae family, were found. Furthermore, two significant proteins, enolase (n=1) and 50S ribosomal protein L7/L12 (n=2), were from Clostridiaceae, whereas the other significant proteins were not clearly correlated with the metagenomics results. It needs to be taken into account that protein databases can be more accurate for certain families than others, and this can affect the taxonomical assignment. These results were represented in the PCA and the HCA (Fig 6AC), where it can be observed that the FMT group clusters closer to the LFD group than the HFD group, indicating that the functionality of gut microbiota after FMT is more similar to that of healthy donor microbiota.

Additionally, a correlation analysis between the metagenomics and metaproteomics results was performed. As shown in Fig 7, the profile that was obtained was very similar in terms of the correlation between the metagenomics and metaproteomics results for the relative abundance of the three significant families that contained all of the significant proteins. These results can be explained by the similarities between these three families, since they are from the same order (Clostridiales) and phylum (Firmicutes). Moreover, clustering of the proteins that belong to the same family or families that are taxonomically related is also observed. Overall, one of the most interesting proteins in our study was glyceraldehyde-3-phosphate dehydrogenase from Clostridiaceae, which has a moderate positive correlation (r=0.5) with the metagenomics results. The slope for this correlation was > 1.3 (metagenomics vs metaproteomics), which indicates that the abundance of this protein and, therefore, its function was diminished in the HFD group vs the LFD and FMT groups. Glyceraldehyde-3-phosphate dehydrogenase was found to control NAD-dependent glycolytic activity in some Clostridium species. It is known that Clostridium species are gram-positive obligate anaerobes and typically perform butyric acid fermentation that is carried out during the exponential growth phase, and this generates acetate and butyrate as the main fermentation products from glucose. In this sense, butyrate products could be considered short-chain fatty acids (SCFAs) produced by gut microbiota [39,40].

**Fig 7.**
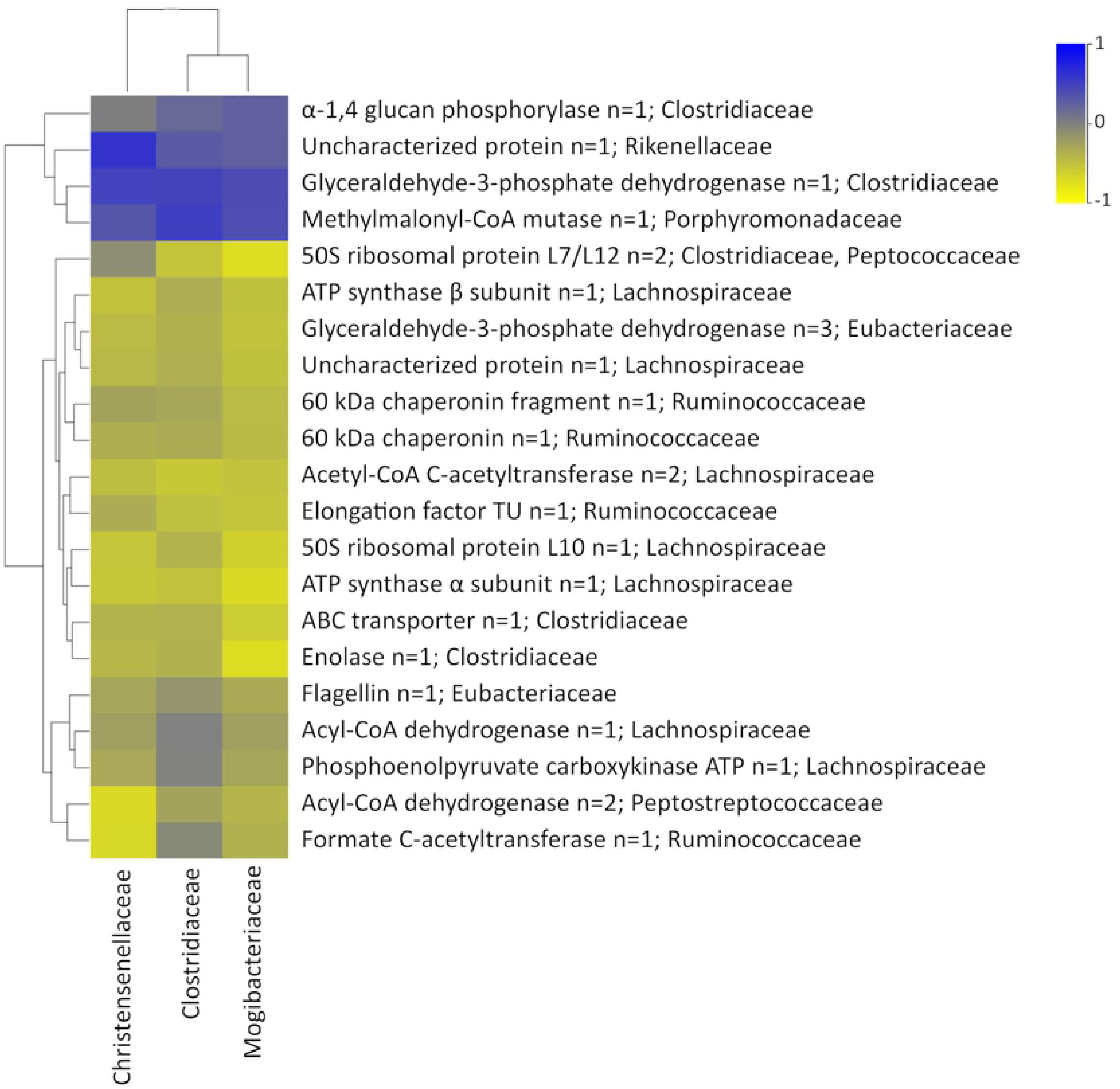
Significant correlations between families and proteins in the LFD, HFD and FMT groups.

Finally, the corroboration of our findings was challenging due to the low number of studies that have been published that have assessed the relationship between metaproteomics and hosts, and even fewer have examined this after FMT. Nevertheless, a study that was performed in rats fed a diet rich in fat and rats fed a chow diet found that Firmicutes were presumed to benefit from a high-fat diet [41], which is contrary to our findings that showed that the abundance of Firmicutes was reduced in the HFD group. However, these differences depend on the part of the colon that is analyzed, as shown in this paper. Moreover, another study performed in pigs found that proteins involved in carbohydrate metabolism showed the most changes in HFD animals, but proteins involved in this process were not changed in our study [42].

## Conclusions

The gut microbiota is essential for maintaining health and has a primary role in metabolism and homeostasis, and its alteration during obesity is a problem that needs to be addressed. Our results applied a combination of metagenomics and metaproteomics approaches to confirm some previous observations: (i) the diet can alter the biochemical composition of the gut microbiota either by shifting the phylotype composition or the activity of bacterial cells; (ii) antibiotics disrupt microbiota biodiversity; (iii) FMT is effective in recolonizing the gut microbiota and in restoring some metabolic functions. When testing these three microbiota modulation strategies, different changes were observed in the bacterial metaproteome, demonstrating that every single change in the host environment can affect microbiota function. In addition to results observed over a short-term period of time [16,18], these findings show that a HFD has a major impact on the mouse cecal microbiota that extends beyond compositional changes to major alterations in bacterial physiology, and FMT can be considered a new strategy to treat obesity.

Moreover, this study reaffirms that metaproteomics should be a complementary tool used along with metagenomics and that combining the results of both approaches can result in the improved characterization of cecal microbiota.

